# A Reusable Non-Adhesive Chest-Wall Acoustic Wearable Estimates Respiratory Rate During Rest and Exercise

**DOI:** 10.64898/2026.06.15.732303

**Authors:** Lisa Haxel, Antonia Schroff, Christopher Fennell, John W. Dickinson

## Abstract

Respiratory rate is a clinically and behaviourally informative signal, yet continuous monitoring outside quiet rest remains difficult. Wearable systems often infer breathing indirectly from cardiovascular or motion surrogates, while direct chest-wall sensing has typically depended on skin adhesion or controlled laboratory conditions. We evaluated Alveos One, a body-coupled acoustic and inertial chest-wall wearable, to test whether a reusable magnet-through-textile form factor can estimate respiratory rate across rest, controlled breathing, motion tasks, and graded exercise. In a single-session protocol, we analysed recordings from 20 healthy adults referenced to structured light plethysmography, breath-by-breath spirometry, and paced-breathing targets. Estimates fell within 2 breaths/min of the reference for 94.6% of valid windows during rest and controlled breathing and 79.3% during exercise; Bland–Altman bias was − 0.24 breaths/min during rest and controlled breathing and −1.07 breaths/min during exercise. Secondary analyses compared the magnet configuration with a skin-adhered patch and assessed airway-mode classification and motion-related operating limits. These findings support reusable, non-adhesive chest-wall acoustic sensing as a practical route to longitudinal respiratory-rate monitoring, and identify rising motion intensity and high ventilatory demand as the principal limits on confident reporting.

## 1 Introduction

Respiratory rate is a basic vital sign and a compact readout of physiological state, yet it is captured less consistently than heart rate, blood pressure, or oxygen saturation. In acute care, abnormal respiratory rate can precede clinical deterioration (1). Outside acute care, breathing rate and breathing pattern change with physical effort, sleep, stress, autonomic state, posture, speech, and voluntary control (2, 3). These properties make breathing attractive for continuous monitoring, but they also make the measurement difficult: a useful wearable must estimate respiratory rate across a wider range of behaviours than quiet seated rest.

Most consumer-compatible wearables do not measure breathing directly. Instead, they infer respiratory rate from respiratory modulation of electrocardiography, photoplethysmography, interbeat intervals, or inertial signals (4–6). These approaches can perform well when the physiological and measurement context is favourable. For example, nocturnal respiratory rate derived from heart-rate-enabled Fitbit data has been validated with sub-breath-per-minute average error in sleep-study data (7); WHOOP respiratory rate showed low bias during polysomnography in healthy adults (8); Samsung Galaxy Watch respiratory-rate estimates showed low overnight error against polysomnography-derived references, although accuracy decreased in severe obstructive sleep apnea (9); and Oura reports a PPG-derived nighttime average respiratory-rate metric for consumer longitudinal tracking (10). These studies show that unobtrusive respiratory-rate monitoring is technically feasible and meaningful to users. They also define the remaining measurement gap: most consumer systems emphasise nocturnal or daily averages inferred from cardiac or wrist-derived surrogates, rather than direct respiratory waveform information during deliberate breathing, posture changes, speech, movement, and exercise.

The waveform distinction matters because respiratory rate is only one summary of breathing. Breath-to-breath variability, inspiratory and expiratory timing, inspiratory:expiratory ratio, duty cycle, and amplitude-like respiratory descriptors are used in respiratory physiology and tidal-breathing analysis (11–14). Audio-based respiratory-monitoring studies have also estimated exhale duration, breathing phase, and inhale-to-exhale ratio from acoustic recordings when breath boundaries are reliable (15, 16). A direct chest-wall acoustic waveform could therefore support richer respiratory phenotyping than an overnight respiratory-rate average alone, provided that waveform-derived features are validated against appropriate references.

Direct respiratory sensing can address that gap, but it introduces its own form-factor problem. Contact-based methods, including strain, impedance, inertial, temperature, and acoustic sensing, preserve more of the respiratory mechanics than indirect cardiac surrogates, but they remain sensitive to placement, coupling, and movement (17). Chest-wall mechano-acoustic sensing offers a complementary route: a body-coupled acoustic sensor records mechanical and airflow-related vibrations close to the respiratory source, while inertial channels help characterise motion and sensor context. Recent chest-wall acoustic systems show promising agreement with clinical references, but performance can degrade during movement, speech, and changes in breathing mode (18).

The practical question is therefore not only whether chest-wall acoustics can estimate respiratory rate, but whether they can do so in a reusable form factor. A skin-adhered patch provides stable coupling to the chest wall, but adhesive wear adds consumables, skin burden, and friction for repeated daily use. A reusable sensor coupled through clothing by magnets could remove these barriers, but only if textile coupling preserves respiratory-rate accuracy during realistic tasks and exercise. The central question in this study is whether such a reusable, non-adhesive configuration can support accurate respiratory-rate monitoring across the operating range relevant to everyday use.

We evaluated the Alveos One chest-wall acoustic wearable in 20 healthy adults during a structured single-session protocol. The primary endpoint was respiratory-rate agreement for the reusable magnet-through-textile form factor against task-appropriate references: structured light plethysmography during unmasked tasks (11, 12), breath-by-breath spirometry during masked and exercise tasks, and instructed rates during paced breathing. A skin-adhered patch was retained as a paired secondary comparator. Additional analyses tested whether the same acoustic signal could recover controlled oral versus nasal breathing (19) and define the conditions under which motion and high ventilatory demand reduce confidence.

## 2 Materials and Methods

### 2.1 Study Design and Participants

We conducted a single-session laboratory validation study in healthy adults (Figure 1). The primary endpoint was agreement between Alveos One respiratory rate and the corresponding laboratory reference for the magnet-through-textile form factor. Secondary endpoints were paired form-factor agreement between the magnet and adhesive configurations, controlled oral/nasal breathing classification, and operating-limit analyses relating respiratory-rate error to signal quality, motion, and ventilatory demand.

**Fig. 1.**
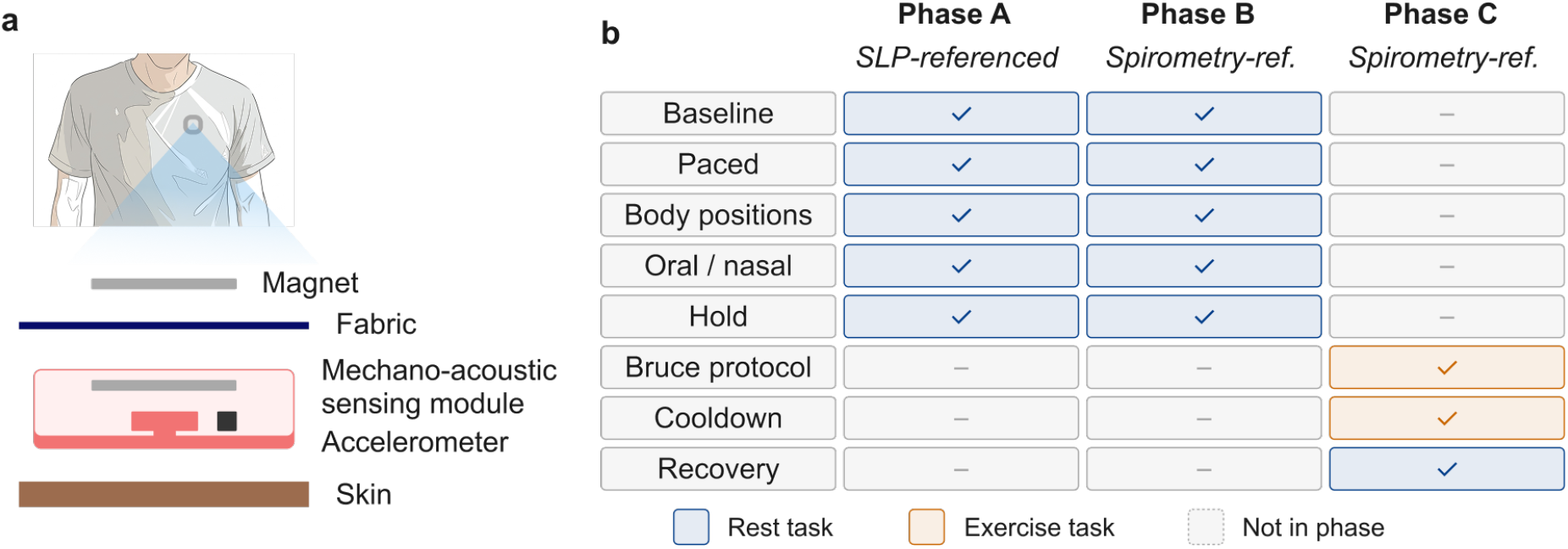
Sensing concept and study design. (**a**) Alveos One records a body-coupled chest-wall acoustic signal and 3-axis acceleration, either as a skin-adhered patch or as a reusable magnet-through-textile sensor coupled across a compression top. (**b**) Protocol overview. Phase A comprised unmasked rest, paced breathing, posture, motion, speech, and controlled oral/nasal breathing tasks; Phase B repeated selected respiratory tasks under spirometry and added short breath holds; Phase C comprised a Bruce-style graded treadmill test followed by cool-down and recovery.

Twenty-four adults (6 female, 18 male; age range 21–64 years) were recruited from the University of Kent community. Exclusion criteria were contraindications to maximal exercise on the Physical Activity Readiness Questionnaire, significant or active respiratory disease, pregnancy, adhesive or chest-strap intolerance, musculoskeletal or balance limitations, resting oxygen saturation below 94% on room air, or resting blood pressure outside the laboratory safety range. Participants were asked to avoid strenuous exercise and alcohol for 24 h, caffeine and nicotine for 3 h, and heavy meals for 2 h before testing; they were also asked to avoid lotions or oils on the upper chest on the testing day. Four participants were excluded from respiratory-rate extraction and oral/nasal classification: three because the Alveos One devices used for recording ran outdated firmware, and one because magnet-form-factor data were unavailable. The final analysis sample therefore comprised 20 participants. All participants gave written informed consent. All participants gave written informed consent.

### 2.2 Device, Form Factors, and Reference Measurements

Alveos One records a body-coupled acoustic signal from the chest wall together with 3-axis acceleration. For each form factor, the acoustic channel was sampled at 8000 Hz and the three accelerometer axes were sampled at 208 Hz. The acoustic channel was recorded using a body-coupled acoustic sensor. All included recordings used Alveos One EVT1 with firmware version v1.1.1.

Each participant wore two devices concurrently to allow a within-participant form-factor comparison. The reusable magnet-through-textile sensor was positioned on the upper anterior chest along the parasternal line, lateral to the sternum and below the sternal angle, corresponding approximately to the region between the second and third intercostal spaces. The sensor was coupled over a fitted compression top and oriented with the charging port facing downwards. The adhesive patch was placed superior to the magnet sensor on the upper chest. In female participants, magnet placement was kept close to the sternum when needed to ensure that the sensor lay flat against the ribcage rather than over soft tissue. The magnet-through-textile configuration was the prespecified primary form factor because it represents the intended reusable everyday-use configuration.

Reference measurements were matched to task constraints. During masked respiratory tasks and graded exercise, a breath-by-breath gas analyser (MetaLyzer 3B; CORTEX Biophysik GmbH, Leipzig, Germany) provided respiratory rate and ventilation measures. During unmasked tasks, structured light plethysmography (Thora-3DI; PneumaScanP2, CS2001, PneumaCare Ltd., Cambridge, UK), operated via PneumaView software (PneumaCare Ltd., Cambridge, UK), provided a thoraco-abdominal respiratory-rate reference without requiring a mask. During paced-breathing tasks, the instructed breathing rate was retained as an additional target reference. At the start of the visit, oxygen saturation was measured using a finger pulse oximeter (PRO-F4; Promise Technology Co., Ltd., China) to confirm resting SpO_2_ above 94% on room air, and blood pressure was measured using an automated blood-pressure monitor (B26; GPZON, Shenzhen Jzmr Technology Co., Ltd., Shenzhen, China).

### 2.3 Protocol and Endpoints

Each participant attended one laboratory visit lasting approximately 2–2.5 h. A trained investigator supervised screening, instrumentation, the recording protocol, device removal, skin inspection, and questionnaires. We generated synchronisation markers at protocol start and at each phase boundary across all recording systems.

The protocol contained three recording phases. Phase A used unmasked SLP-referenced tasks: quiet baseline, speech, scripted motion artefacts, paced breathing at 6, 8, 12, 15, and 20 breaths/min, seated/standing/supine postures, and controlled oral-only and nasal-only breathing. Phase B used spirometry-referenced masked tasks: quiet baseline, paced breathing, short voluntary breath holds, posture variation, and masked airway tasks where equipment allowed. Phase C was a Bruce-style graded treadmill test continued until volitional fatigue, a predefined safety endpoint, participant request, or investigator termination, followed by cool-down and recovery. Supplementary Table 1 summarises the task-to-reference mapping used for analysis.

The primary respiratory-rate endpoint was the percentage of valid magnet-form-factor estimates within 2 breaths/min of the reference, complemented by Bland–Altman bias and 95% limits of agreement (20). Secondary respiratory summaries included the mean absolute error, root mean squared error, mean absolute percentage error, Lin’s concordance correlation coefficient, proportional bias, and valid-window fraction. The same agreement framework was used for the adhesive comparator.

### 2.4 Respiratory-Rate Estimation and Reference Pairing

All Alveos One recordings were processed offline. Acoustic and reference streams were aligned with the protocol synchronisation markers and checked against salient task events. A fixed 3.0 s edge interval was removed from the beginning and end of each Alveos One recording to reduce handling and filter-settling artefacts. Recordings shorter than 11.0 s before trimming were excluded at the loading stage; endpoint analyses further required at least one valid paired Alveos One–reference respiratory-rate estimate.

Respiratory rate was estimated from the acoustic channel after downsampling to 100 Hz, applying Gaussian smoothing and polynomial detrending. We used task-specific sliding windows, with shorter steps for rest and controlled-breathing tasks and longer steps for exercise tasks. Offline zero-phase Butterworth filtering was used for this validation analysis. To account for movement-dependent waveform changes, accelerometer-derived motion was used to assign each recording to a low-, moderate-, or high-motion regime, and a predefined respiratory band-pass filter was selected for that regime. The processed respiratory waveform was then analysed for repeated extrema; peak-based and trough-based rate estimates were combined into one respiratory-rate estimate per window. Estimates outside 3–50 breaths/min were excluded.

A respiratory-rate signal-quality score (RR–SQS) was computed for each retained window from waveform complexity, periodicity, and interval regularity. Windows below the locked RR–SQS threshold were masked before agreement analysis. Bounded interior gaps between valid respiratory-rate estimates were linearly interpolated, whereas low-quality leading or trailing windows remained missing. The full parameter set, including window lengths, step sizes, filter definitions, edge-artefact handling, interpolation rules, is provided in the GitHub repository **alveoslabs/alveos-respiratory-validation**. The RR–SQS definition can be found in Section S1.4.

For spirometry-referenced tasks, each Alveos One window was paired with the median spirometry breathing frequency over the same temporal window. For SLP-referenced tasks, the mean valid Alveos One estimate for each task was paired with the corresponding task-level SLP respiratory-rate estimate. For paced-breathing tasks, the instructed target rate was retained as an additional reference. We retained a paired observation only when both the Alveos One estimate and the reference value were finite after quality masking and interpolation.

### 2.5 Secondary Analyses

The adhesive and magnet-through-textile form factors were compared within participant and task wherever both recordings were valid. The paired comparison used participant-level differences in mean absolute error, Bland–Altman bias, limits-of-agreement width, valid-window fraction, and percentage within 2 breaths/min. Rest/controlled and exercise conditions were summarised separately because exercise imposes the strongest combined challenge from motion, sweat, ventilation, and textile coupling.

Controlled oral-only and nasal-only Phase A blocks tested whether the magnet-form-factor acoustic signal contained airway-mode information. The prespecified endpoint used segment-level acoustic envelope features, minimum Redundancy–Maximum Relevance (mRMR) feature selection, robust scaling, and a shrinkage linear discriminant classifier. The primary classification strategy was participant-specific calibration with 20 s per class and evaluation on temporally separated held-out segments. Feature selection, scaling, and classifier fitting were repeated within each calibration draw using only calibration data before evaluation on held-out segments. Balanced accuracy was the primary metric; area under the receiver operating characteristic curve, sensitivity, specificity, confusion matrices, recording-level metrics, confidence-thresholded performance, zero-shot transfer, source-plus-target training, and segment-duration sensitivity were secondary analyses.

### 2.6 Statistical Analysis and Data Availability

Continuous variables are reported as mean ± standard deviation or median (IQR), according to distributional shape. Categorical variables are reported as counts and percentages. Bland–Altman analyses summarised agreement, and proportional bias was assessed by regressing paired differences against pair means. Window-level observations were used for agreement plots, descriptive summaries, and operating-limit analyses. Because overlapping windows from the same participant are correlated, cohort-level summaries and inferential comparisons were based on participant-level estimates. Confidence intervals for participant-level summaries and oral/nasal classification were estimated as percentile bootstrap intervals using 10,000 resamples across participants. For oral/nasal classification, repeated calibration draws were averaged within participant before resampling. For paired form-factor comparisons, differences between magnet and adhesive estimates were computed within participant before cohort-level summarisation.

## 3 Results

### 3.1 The magnet form factor estimates respiratory rate accurately across rest and exercise

We first tested whether the reusable magnet-through-textile configuration could estimate respiratory rate across the main validation conditions. During rest and controlled-breathing tasks, 94.6% of valid estimates fell within 2 breaths/min of the reference, with a Bland–Altman bias of -0.24 breaths/min and a 95% limits-of-agreement width of 4.71 breaths/min (Figure 2a,c,d). During graded exercise, where ventilation, movement, sweat, and textile coupling were expected to challenge the measurement, 79.3% of valid estimates remained within 2 breaths/min, with a bias of -1.07 breaths/min and a 95% limits-of-agreement width of 4.80 breaths/min (Figure 2b–d). Across participants, the median percentage within 2 breaths/min was 91.5% (IQR 86.3–94.9%; Figure 2c). Proportional bias was statistically detectable but small in magnitude (slope -0.05, *p* < 0.001), indicating only a minor systematic shift across the respiratory-rate range. These results show that the magnet configuration preserved respiratory-rate agreement from rest through exercise, with lower but still substantial threshold accuracy under high ventilatory and movement demand.

**Fig. 2.**
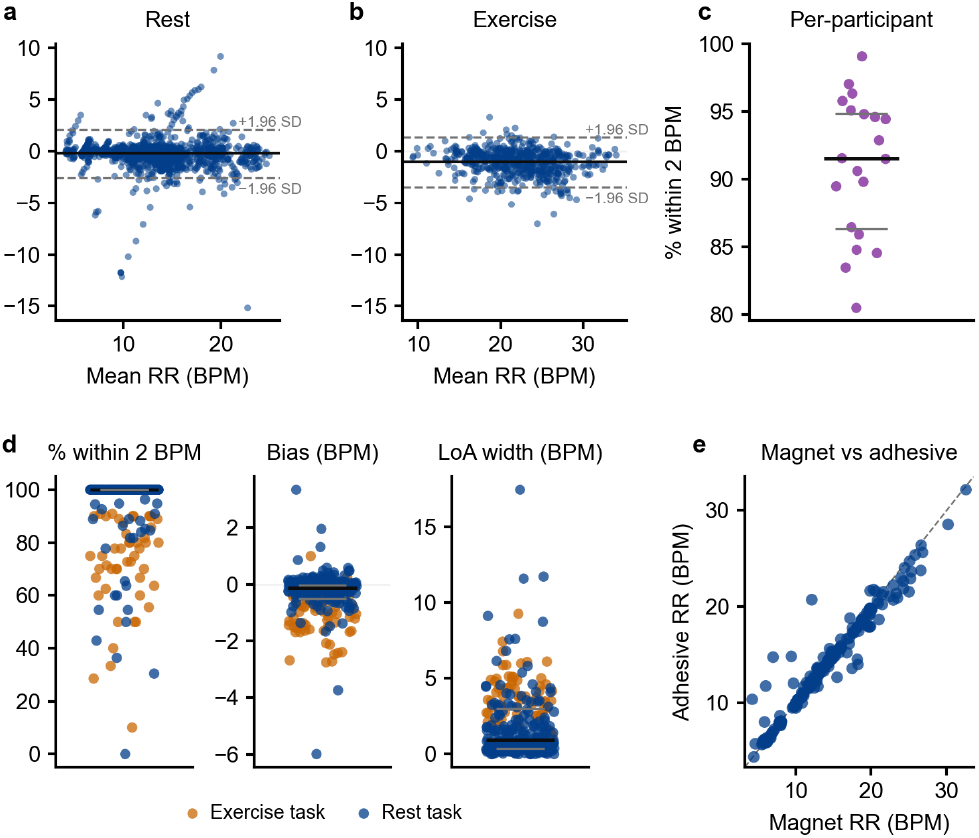
Respiratory-rate validation for the magnet-through-textile form factor. (**a**) Bland–Altman agreement for rest and controlled-breathing tasks against the combined reference (bias -0.24 breaths/min; 95% LoA-width 4.71 breaths/min; 3400 paired observations from 20 participants). (**b**) Bland–Altman agreement for graded exercise (bias -1.07 breaths/min; 95% LoA-width 4.80 breaths/min; 670 paired observations from 20 participants). (**c**) Participant-level percentage within 2 breaths/min across rest/controlled and exercise conditions. (**d**) Recording-level percentage within 2 breaths/min, bias, and limits-of-agreement width. Black horizontal lines in (**c**,**d**) indicate the median; grey lines indicate the IQR. (**e**) Magnet versus adhesive form-factor agreement: window-level respiratory-rate estimates from the magnet sensor plotted against simultaneous estimates from the adhesive patch, pooled across all rest and exercise tasks. The dashed line indicates identity.

### 3.2 The reusable magnet form factor preserves agreement relative to the adhesive patch

Replacing direct skin adhesion with textile coupling could reduce mechanical coupling and degrade respiratory-rate agreement. We therefore compared simultaneous magnet and adhesive recordings to quantify this form-factor trade-off (Figure 2e). Across rest and controlled-breathing tasks, the magnet and adhesive configurations performed similarly. Relative to the adhesive patch, the magnet configuration reduced mean absolute error by 0.08 breaths/min and increased the percentage of estimates within 2 breaths/min by 0.5 percentage points. Bland–Altman agreement was also comparable between form factors at rest and during controlled breathing, with similar bias (-0.24 versus -0.27 breaths/min) and limits-of-agreement width (4.72 versus 4.72 breaths/min).

During exercise, the reusable configuration did not show the expected loss of agreement. Relative to the adhesive patch, the magnet configuration reduced mean absolute error by 0.66 breaths/min and increased the percentage of estimates within 2 breaths/min by 14.3 percentage points. Bias was smaller for the magnet configuration than for the adhesive patch (-1.07 versus -1.78 breaths/min), and the limits-of-agreement width was narrower (4.80 versus 7.02 breaths/min). Under the tested laboratory conditions, the magnet-through-textile configuration therefore preserved respiratory-rate agreement relative to direct skin adhesion and performed at least comparably during exercise, where form-factor differences were expected to matter most.

### 3.3 Motion and ventilatory demand define the respiratory-rate operating envelope

We next examined when respiratory-rate estimation degraded. Error was concentrated in windows with low RR–SQS and in recordings with larger movement or larger respiratory-rate excursions (Figure 3). Above the locked RR–SQS threshold, signed error remained centred near zero. In contrast, low-quality windows occurred most often during speech, motion, and posture transitions, where the waveform no longer supported a stable respiratory-rate estimate (Figure 3a). At the recording level, median absolute error increased from 0.13–0.14 breaths/min in recordings with low accelerometer or respiratory-rate range to 0.57–0.62 breaths/min in recordings with high accelerometer or respiratory-rate range (Figure 3b). Visual inspection of low-quality examples identified three recurring failure modes: motion contamination, breath-hold or task-transition artefacts, and irregular breathing with transient large excursions (Figure 3c). Together, these analyses identify movement and rapid changes in breathing pattern as the main operating limits, rather than quiet posture or regular resting respiration.

**Fig. 3.**
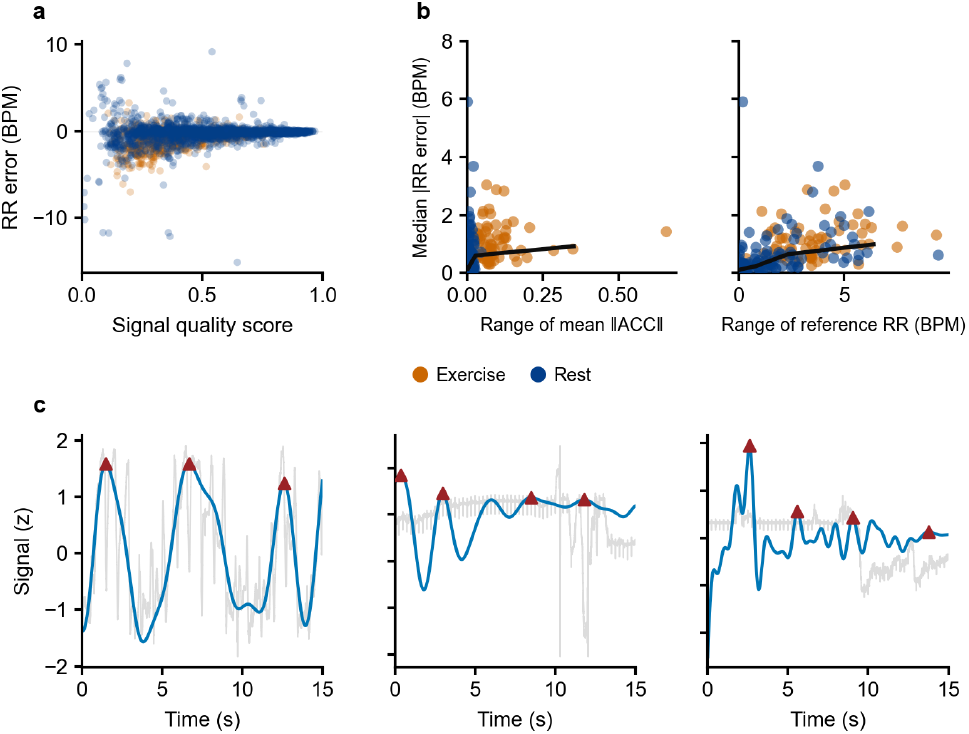
Operating limits of respiratory-rate estimation. (**a**) Window-level respiratory-rate error (Alveos One minus reference) as a function of RR–SQS, stratified by rest/controlled and exercise conditions. (**b**) Recording-level median absolute error against the within-recording range of accelerometer magnitude and the within-recording range of reference respiratory rate. (**c**) Representative low-quality windows illustrating motion contamination, breath-hold or transition-related amplitude change, and irregular breathing with a transient large excursion.

### 3.4 Chest-wall acoustics support calibrated oral/nasal airway-mode classification

Respiratory rate captures only one aspect of the chest-wall acoustic signal. We therefore tested whether controlled oral and nasal breathing could also be distinguished from the same recordings. Using controlled oral-only and nasal-only breathing blocks, we trained segment-level classifiers on short participant-specific calibration segments and evaluated them on temporally separated held-out data (Figure 4a).

**Fig. 4.**
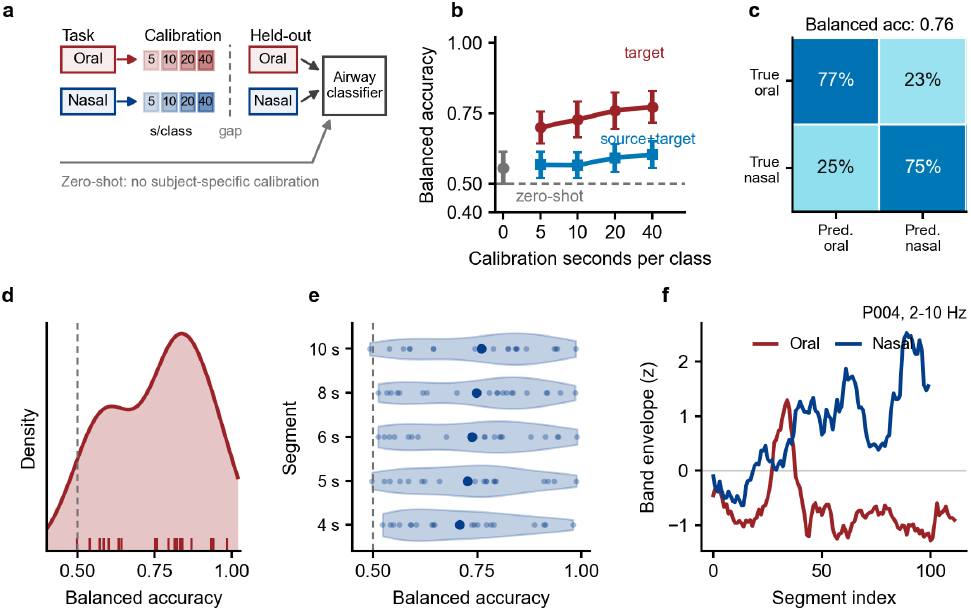
Oral-versus nasal-breathing classification from chest-wall acoustic features. **(a)** Participant-specific calibration design using controlled oral-only and nasal-only recordings with temporally separated held-out test segments. (**b**) Calibration-duration curve for segment-local classification without recording-context features, comparing zero-shot transfer, target-only participant-specific calibration, and source-plus-target training. Points and error bars summarize participant-level performance after repeated calibration draws were averaged within participant. (**c**) Participant-balanced row-normalised confusion matrix for the target-only end-point using 20 s/class calibration, 10 s segments, 1 s step, mRMR feature selection, and a shrinkage linear discriminant classifier. (**d**) Distribution of participant-level balanced accuracy for the locked 20 s/class target-only endpoint. (**e**) Segment-duration sensitivity for the same target-only calibration setting, comparing 4, 5, 6, 8, and 10 s segments. (**f**) Representative participant-level 2–10 Hz band-envelope trajectories during controlled oral and nasal breathing. All analyses used Phase A controlled oral- and nasal-breathing blocks and evaluated held-out segments separated from calibration data.

**Fig. 5.**
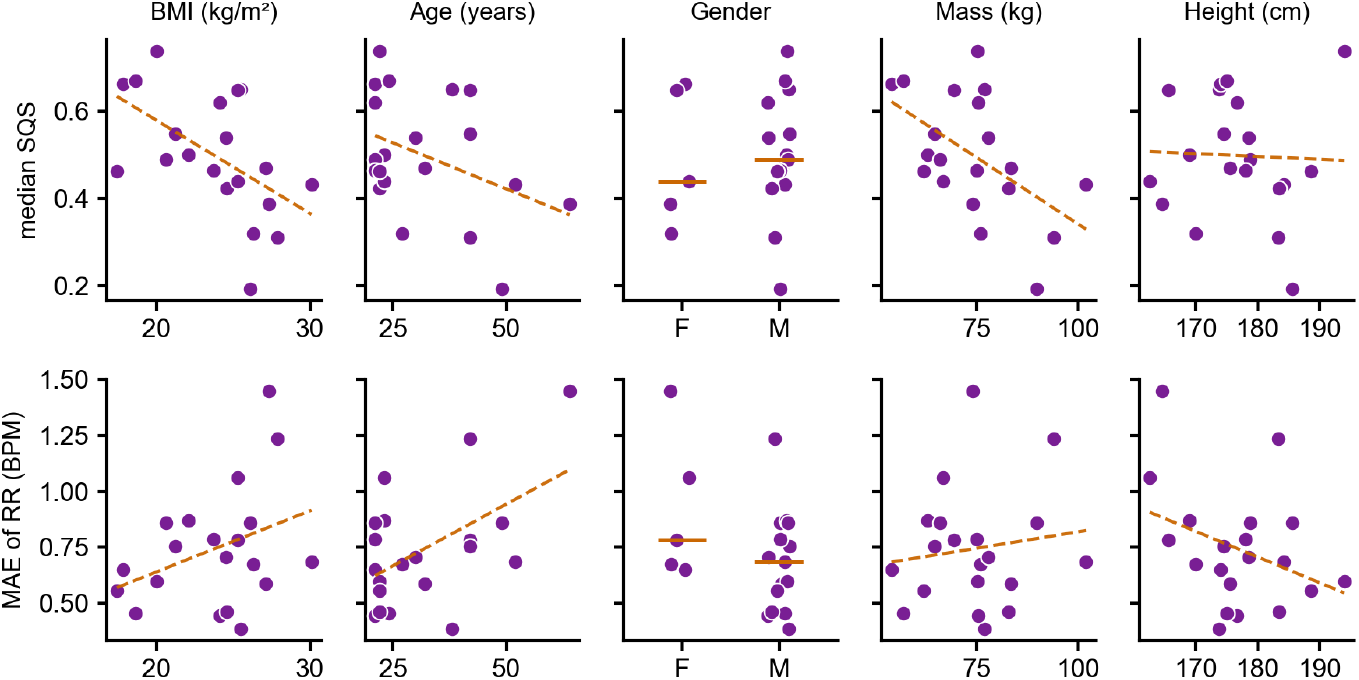
Per-participant median RR–SQS (top) and respiratory-rate MAE (bottom) against BMI, age, gender, body mass, and height. Orange lines are ordinary least-squares fits (continuous variables) or group medians (gender); reported associations use Spearman’s *ρ*. Exploratory, *n* = 20.

**Fig. 6.**
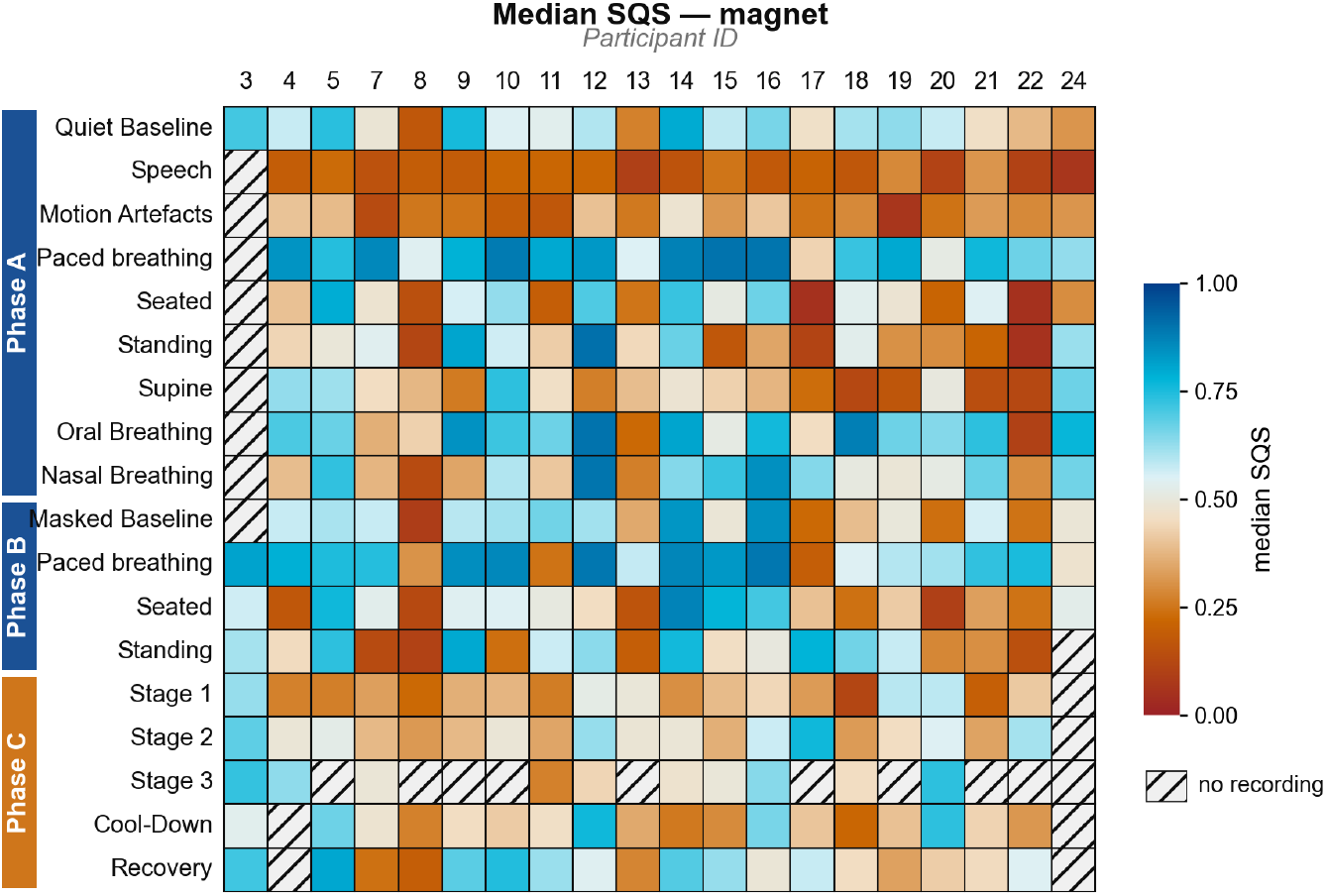
Median RR–SQS for the magnet-through-textile form factor by task (rows, grouped by phase) and participant (columns). Hatched cells indicate tasks with no valid recording. Colour scale runs from low (0) to high (1) median signal quality.

**Fig. 7.**
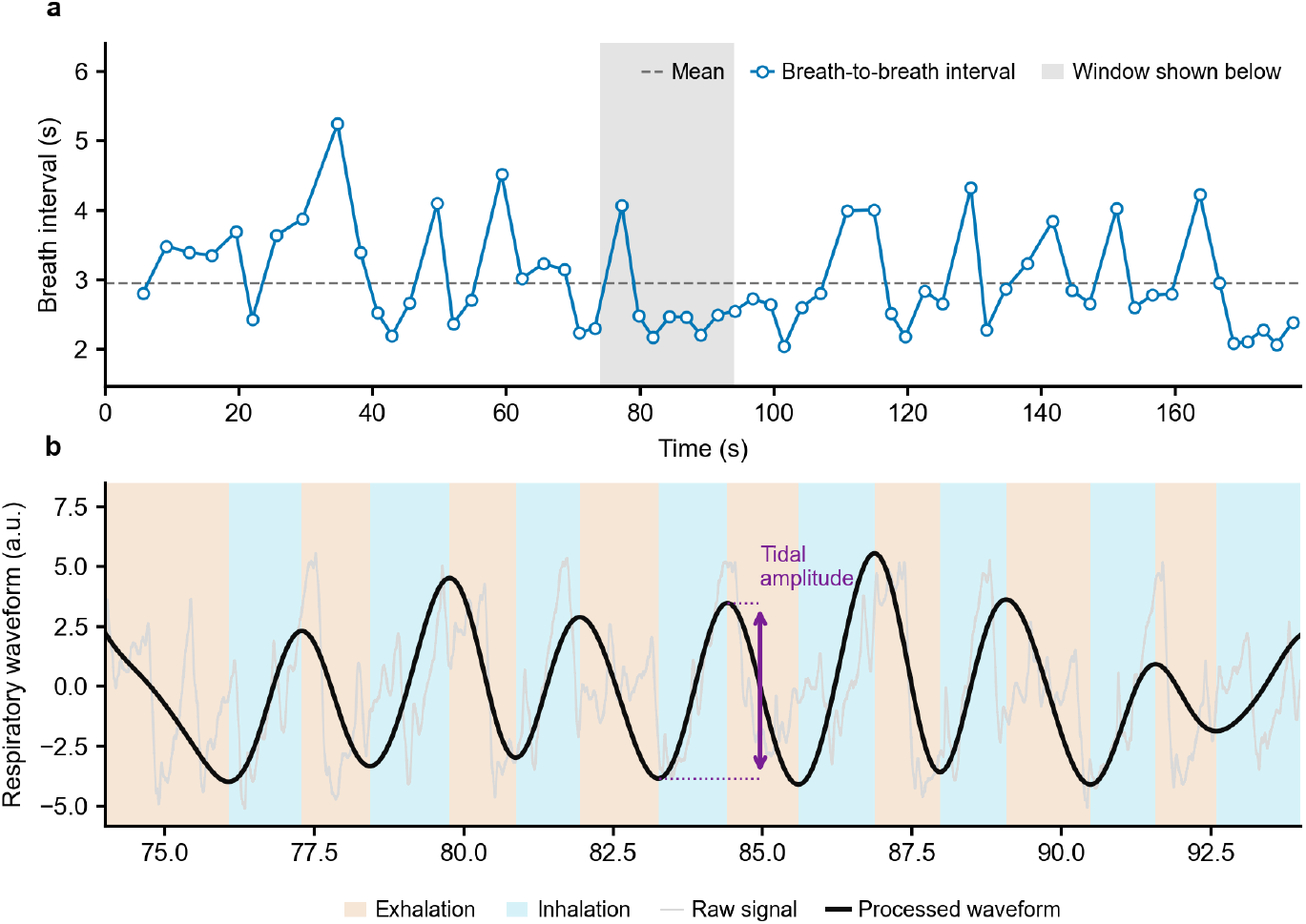
Exploratory waveform-derived respiratory descriptors from the magnet form factor (representative recording from Stage 1 of the Bruce protocol). (**a**) Breath-to-breath interval time series; the dashed line marks the mean interval and the shaded span indicates the window detailed in (**b**). (**b**) Raw acoustic signal (grey) and processed respiratory waveform (black) over the shaded window, with detected inhalation and exhalation phases shaded and the peak-to-trough tidal-amplitude proxy annotated for one breath. Descriptors shown are illustrative and were not validated against an independent reference (see Section 4).

**Fig. 8.**
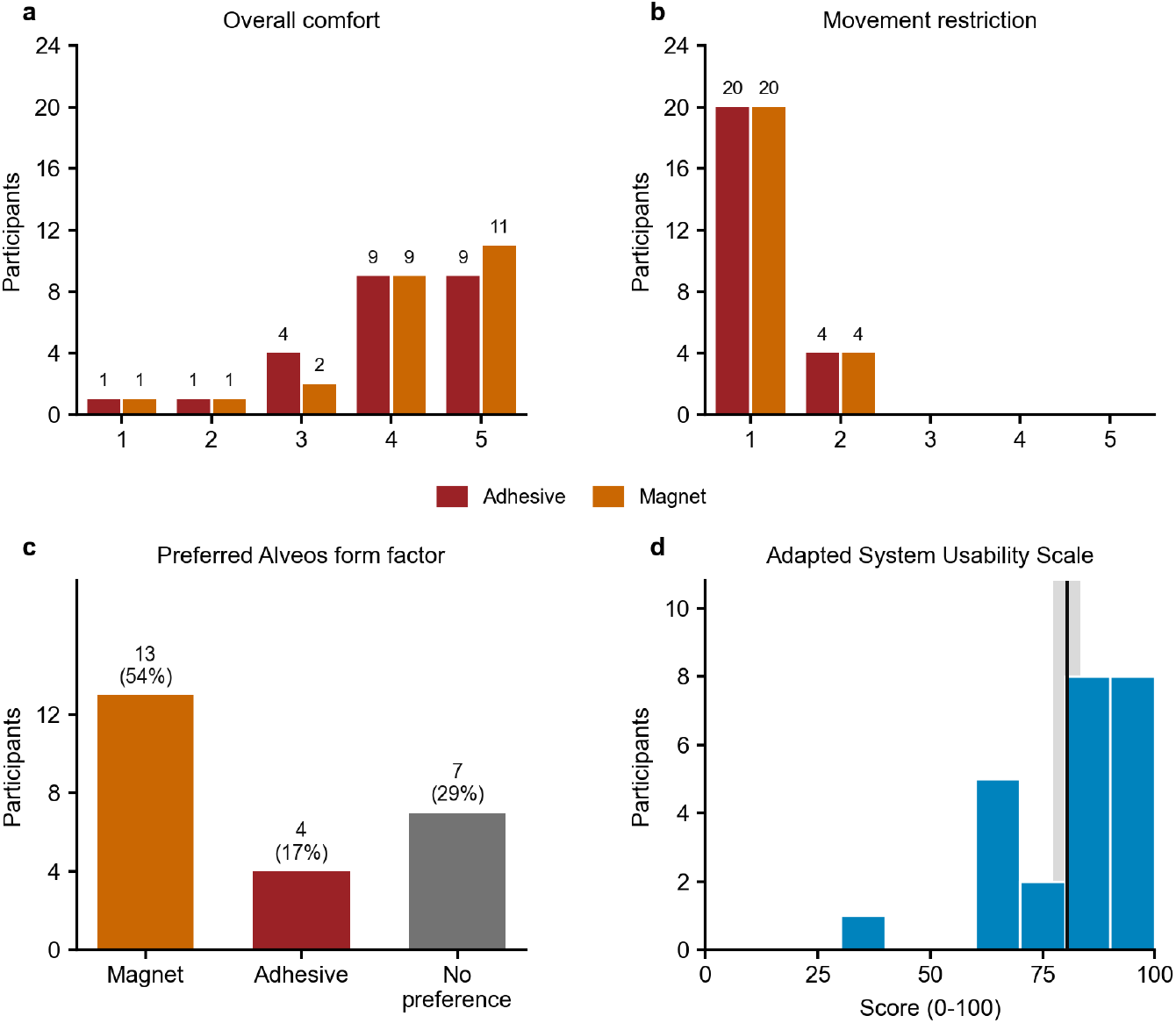
Participant-reported acceptability of the two Alveos One form factors (*n* = 24). (**a**) Overall comfort rating per device (1 = very uncomfortable, 5 = very comfortable). (**b**) Movement restriction during the exercise test (1 = not at all, 5 = significantly restricted). (**c**) Preferred form factor for hypothetical extended daily or overnight use. (**d**) Distribution of adapted System Usability Scale scores for the Alveos One system overall (0–100); the vertical line marks the median and the shaded band the inter-quartile range.

With 20 s/class of participant-specific target-only calibration, the classifier reached a balanced accuracy of 0.76 (95% CI 0.70–0.82; Figure 4b–d). Overall accuracy was 0.75 and AU-ROC was 0.78. The participant-balanced confusion matrix showed similar performance for both classes, with 77% correct classification of oral-breathing segments and 75% correct classification of nasal-breathing segments (Figure 4c). For statistical inference, repeated calibration draws were first averaged within participant, and the three planned oral/nasal tests were corrected using the Holm method. The resulting participant-level balanced accuracies were above the chance level of 0.5 (one-sample t-test: *t*(19) = 7.79, Holm-adjusted *p* = 7.46 ×10^−7^).

Performance depended strongly on participant-specific calibration. Target-only balanced accuracy increased from 0.70 with 5 s/class calibration to 0.73 with 10 s/class, 0.76 with 20 s/class, and 0.77 with 40 s/class. In contrast, zero-shot transfer without participant-specific calibration remained near chance, reaching 0.56 balanced accuracy. Adding source-participant data did not improve performance over target-only calibration: source-plus-target training reached 0.59 at 20 s/class and 0.60 at 40 s/class. At the locked 20 s/class endpoint, target-only calibration outperformed both zero-shot transfer (mean paired difference 0.20; paired t-test: *t*(19) = 5.98, Holm-adjusted *p* = 1.86 × 10^−5^) and source-plus-target training at the same calibration duration (mean paired difference 0.17; paired t-test: *t*(19) = 5.82, Holm-adjusted *p* = 1.86 × 10^−5^). Oral/nasal discrimination was therefore primarily a calibrated within-participant read-out rather than a robust zero-shot transfer task.

Longer analysis segments provided more stable airway-mode information. In the target-only calibration setting, balanced accuracy increased monotonically across the tested segment durations, from 0.71 for 4 s segments to 0.73 for 5 s, 0.74 for 6 s, 0.75 for 8 s, and 0.76 for 10 s segments (Figure 4e). Together, these results suggest that the chest-wall acoustic recordings contain airway-mode information, but that the readout currently depends on short participant-specific calibration and benefits from several seconds of respiratory context.

## 4 Discussion

This study tested whether a reusable, non-adhesive chest-wall acoustic wearable can preserve respiratory-rate agreement across conditions that typically challenge wearable respiratory monitoring. The magnet-through-textile configuration recovered respiratory rate across quiet breathing, controlled breathing, posture changes, speech, motion tasks, and graded exercise. Agreement was strongest during rest and controlled breathing and lower during exercise, where motion, high ventilatory demand, sweat, and changes in textile coupling can distort the respiratory waveform. Paired adhesive recordings tested the form-factor trade-off directly, and controlled oral/nasal recordings showed that the same acoustic signal also contained airway-mode information. The central contribution is therefore not only a respiratory-rate algorithm, but evidence that reusable chest-wall acoustic coupling can support direct, quality-aware respiratory monitoring across active physiological states.

Consumer respiratory-rate metrics provide the relevant benchmark, but they answer a different measurement question. Fitbit, WHOOP, Samsung Galaxy Watch, and Oura studies show that respiratory rate can be estimated from cardiac or wrist-derived signals during sleep, overnight rest, or daily monitoring (7–10, 21). These systems have helped make respiratory rate a practical longitudinal health metric. Their strongest use case, however, is usually an aggregated nocturnal or daily trend. The present study instead tested a direct chest-wall signal during deliberate breathing, posture changes, speech, movement, and exercise, where respiration changes rapidly and where cardiac-surrogate approaches are harder to interpret because heart rate, perfusion, motion, and respiratory drive change together (4, 22, 23). The relevant distinction is therefore not consumer versus research-grade monitoring, but indirect aggregated trend estimation versus direct respiratory waveform sensing in active conditions.

The direct waveform is important because respiratory rate discards much of the respiratory pattern. Once individual breaths can be segmented reliably, the same waveform can define breath-to-breath respiratory-rate variability, inspiratory time, expiratory time, total breath duration, inspiratory:expiratory ratio, inspiratory duty cycle, and amplitude-like descriptors (11–14). Acoustic respiratory recordings and tracheal-sound systems have similarly been used to extract respiratory timing, breathing phase, respiratory rate, and inhale-to-exhale structure (15, 16, 24, 25). These features could matter for applications in exertion monitoring, autonomic biofeedback, sleep phenotyping, respiratory disease monitoring, and behavioural-state inference. In the present study, however, respiratory rate was the validated endpoint. Waveform-derived timing and amplitude features should therefore be presented as candidate outputs for future validation, not as established measurements of flow, volume, or airway mechanics (Supplementary Section S2.3, Supplementary Figure 7).

Signal-quality masking should be interpreted as part of the measurement model rather than as post hoc data cleaning. Respiratory waveform algorithms commonly fail when breath morphology, periodicity, or interval regularity degrades, and signal-quality indices have been proposed to identify unreliable respiratory segments before estimating rate (26, 27). The RR–SQS used here follows the same principle: it allows the system to abstain when the waveform no longer supports a confident estimate. This choice makes accuracy conditional on coverage, so agreement statistics should always be interpreted together with valid-window fraction and the distribution of masked periods. For a deployable system, quality-aware abstention is preferable to always reporting a numerically precise but physiologically unreliable respiratory rate.

The speech task deserves separate interpretation because speech is not simply a motion artefact. Speaking reorganises respiration around phrase planning, phonation, and communicative demand; breathing pauses occur at linguistically meaningful locations, and verbal or cognitively demanding tasks can alter respiratory timing, airflow, and ventilation (28–30). Speech also injects high-energy acoustic and mechanical activity into the same body-coupled signal used for respiratory monitoring. A respiratory-rate detector tuned for repeated tidal-breathing extrema will therefore encounter two problems during speech: the breathing pattern itself becomes less periodic, and the acoustic channel contains non-respiratory voiced energy. The low RR–SQS observed during speech is therefore physiologically and instrumentally plausible, not merely an algorithmic defect.

Speech should be handled as a distinct operating state in future versions of the pipeline. A practical algorithm could first detect speech or phonation from acoustic features, inertial context, or a dedicated speech-activity classifier, then down-weight or suppress standard respiratory-rate estimates during voiced segments. Reporting could resume after a short stable-breathing interval, with the gap marked as low-coverage rather than imputed. A speech-specific pathway could instead estimate phrase-level breathing, pre-utterance inhalations, post-utterance recovery, or speech-breath coordination, working from the wider-band signal and against references suited to speech rather than resting breathing.

The form-factor comparison is the practical hinge of the study. Adhesive patches maximise mechanical coupling, but they add consumables, skin burden, and friction for repeated wear. The magnet-through-textile configuration targets the use case that matters for longitudinal monitoring: a sensor that can be reused, repositioned, and worn without direct skin adhesion. The paired adhesive comparison therefore asks whether this practical form factor pays an unacceptable accuracy penalty. In the present supervised protocol, the reusable configuration preserved respiratory-rate information across rest and exercise sufficiently to justify the next validation step: multi-session home and field studies that quantify adherence, comfort, placement repeatability, garment effects, and data coverage.

Controlled oral-only and nasal-only recordings showed that the magnet-form-factor acoustic signal contains information beyond respiratory rate. With 20 s/class of participant-specific calibration, a segment-local model classified held-out oral and nasal breathing above chance, whereas zero-shot transfer remained near chance. This pattern suggests that airway-mode information exists in the signal but depends on individual coupling, anatomy, or calibration. Future work should test whether calibration can be shortened, whether signatures remain stable across days and garments, and whether models trained during controlled breathing generalise to natural mixed breathing, speech, sleep, and exercise.

Several limitations bound the interpretation. Participants were healthy adults tested in a single supervised laboratory session, so performance may not generalise to older adults, respiratory disease, sleep, high body-mass-index ranges, free-living routines, or prolonged wear. The respiratory-rate pipeline used offline zero-phase filtering; real-time deployment will require causal filtering, latency quantification, and prospective locking of all signal-quality rules. The study also used task-specific references, which was necessary because structured light plethysmography, spirometry, and paced-breathing targets suit different conditions, but this heterogeneity limits simple comparison across the full protocol. Finally, the acoustic waveform should not yet be interpreted as validated breath-by-breath flow or volume. Respiratory rate was the primary endpoint; inspiratory timing, expiratory timing, respiratory variability, airway mode, and amplitude-like descriptors remain secondary or exploratory until validated against appropriate references.

Within these limits, the study advances wearable respiratory monitoring by shifting the emphasis from passive nocturnal trend estimation to direct, reusable, quality-aware chest-wall sensing. Consumer sleep wearables show that respiratory rate can be a robust longitudinal health signal. Alveos One addresses a complementary use case: preserving respiratory information when users breathe deliberately, move, speak, change posture, and exercise. The next translational step is prospective free-living validation in the populations and contexts where continuous respiratory monitoring could change clinical, training, or behavioural decisions.

## 5 Conclusions

A reusable magnet-through-textile chest-wall acoustic wearable estimated respiratory rate accurately across rest, controlled breathing, and graded exercise in healthy adults. Secondary analyses compared the reusable form factor with an adhesive patch and showed that the same acoustic recordings can support airway-mode readouts. These findings support direct chest-wall acoustic sensing for longitudinal respiratory monitoring, with free-living and clinical-population validation as the next priorities.

## Ethics statement

The study was conducted in accordance with the Declaration of Helsinki and approved by the School of Sport, Exercise and Rehabilitation Sciences Research Ethics and Advisory Group (REAG), University of Kent (protocol code 01_2026, approval date 26 January 2026).

## Data availability

Analysis code, figure-generation scripts, and configuration files are available at **alveoslabs/alveos-respiratory-validation**. De-identified data supporting the reported results are available from the corresponding authors on reasonable request, subject to Regulation (EU) 2016/679.

## AUTHOR CONTRIBUTIONS

Conceptualization: L.H. and J.W.D.; methodology: L.H. and A.S.; software: L.H. and A.S.; validation: L.H. and A.S.; formal analysis: L.H. and A.S.; investigation: C.F.; resources: J.W.D; data curation: C.F, L.H.; writing—original draft preparation:

L.H. and A.S.; writing—review and editing: C.F and J.W.D. ; visualization: L.H. and A.S.; supervision: J.W.D. ; project administration: L.H. and J.W.D.; funding acquisition: J.W.D.. All authors have read and agreed to the published version of the manuscript.

## ACKNOWLEDGEMENTS

This research was funded by Alveos Oy.

## COMPETING FINANCIAL INTERESTS

A.S. and L.H. are employees of Alveos Oy, which developed the device under evaluation and funded this study; as such, the funder was involved in the work through their contributions, as specified in the Author Contributions section. C.F. and J.W.D. declare no conflicts of interest.

## S1 Supplementary Methods

### S1.1 Task and Reference Mapping

Tasks were grouped by measurement context. Phase A comprised unmasked tasks referenced to structured light plethysmography and provided the main controlled oral/nasal recordings. Phase B used spirometry-referenced masked respiratory tasks and breath holds. Phase C used spirometry during graded exercise, cool-down, and recovery. Supplementary Table 1 lists the analysis groups, target rates where applicable, and primary reference for each group.

**Table 1.** Task groups and reference modalities used for analysis. Internal recording labels are defined in the released configuration.

| Phase | Task group | Target rate | Primary reference or endpoint role |
| --- | --- | --- | --- |
| A | Quiet baseline | – | SLP respiratory rate |
| A | Speech | – | SLP respiratory rate; quality/artefact analysis |
| A | Scripted motion artefacts | – | Motion and quality analysis |
| A | Paced breathing | 6, 8, 12, 15, 20 breaths/min | SLP and instructed target rate |
| A | Seated, standing, and supine posture | – | SLP respiratory rate; posture/quality analysis |
| A | Controlled oral-only breathing | – | Oral label; SLP respiratory rate |
| A | Controlled nasal-only breathing | – | Nasal label; SLP respiratory rate |
| B | Masked quiet baseline | – | Spirometry respiratory rate |
| B | Masked paced breathing | 6, 8, 12, 15, 20 breaths/min | Spirometry and instructed target rate |
| B | Voluntary breath holds | 15, 20, 25 s | Spirometry flow cessation |
| B | Masked posture tasks | – | Spirometry respiratory rate; posture analysis |
| B | Masked airway task | – | Spirometry respiratory rate; exploratory airway context |
| C | Bruce graded exercise stages | – | Spirometry respiratory rate and ventilation |
| C | Cool-down and recovery | – | Spirometry respiratory rate and recovery context |

### S1.2 Respiratory Preprocessing and Rate Extraction

Alveos One recordings were processed separately by participant, task, and form factor. The acoustic channel was downsampled for respiratory analysis. The three accelerometer axes were converted to vector magnitude, high-pass filtered to suppress gravity, and summarised with a rolling root-mean-square statistic. This motion summary assigned each recording to a low-, moderate-, or high-motion regime, which selected the respiratory band-pass filter. Filters were applied as zero-phase forward–backward filters for offline validation.

Within each sliding window, the processed acoustic waveform was analysed for respiratory extrema. Peak-based and trough-based rate estimates were averaged:

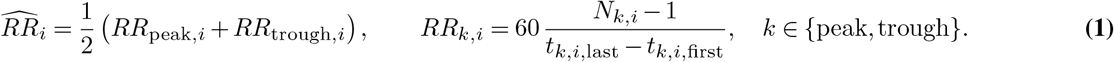

Windows with fewer than two valid peaks and fewer than two valid troughs, or with a final estimate outside 3–50 breaths/min, were excluded. The same extrema supported exploratory descriptors of inspiratory time, expiratory time, breath duration, inspiratory/expiratory ratio, duty cycle, and an acoustic tidal-amplitude proxy. Design-relevant parameters are listed in Supplementary Table 2; remaining tuning constants are specified in the released configuration.

### S1.3 Reference Pairing and Agreement Estimators

For spirometry-referenced tasks, each Alveos One window centred at *t*_*i*_ was paired with the median spirometry breathing frequency in the same temporal window:

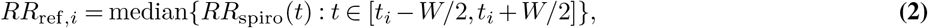

where *W* is the active respiratory-rate window length. For SLP-referenced tasks, the mean valid Alveos One estimate for the task was paired with the corresponding SLP task-level respiratory rate. For paced-breathing tasks, instructed target rate was analysed separately as an additional reference.

For each valid pair, the difference was 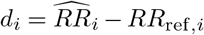 and the pair mean was 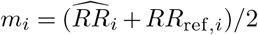. Bland–Altman bias was 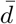 and 95% limits of agreement were 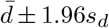 Proportional bias was assessed by regressing *d*_*i*_ on *m*_*i*_. Mean absolute error, root mean squared error, mean absolute percentage error, Lin’s concordance correlation coefficient, and threshold accuracy used standard definitions. For pooled all-reference summaries, paired values were first collapsed to one Alveos One–reference pair per recording and reference source so that long or densely windowed recordings did not dominate the summary.

**Table 2.** Design-relevant respiratory preprocessing and breath-detection parameters. Fine tuning constants are specified in the released configuration.

| Parameter | Value | Description |
| --- | --- | --- |
| Acoustic downsampling rate | 100 Hz | Respiratory waveform analysis. |
| Accelerometer downsampling rate | 100 Hz | Motion summary. |
| Accelerometer high-pass filter | 0.5 Hz | Suppresses gravity before motion estimation. |
| Motion RMS window | 5 s | Window for dynamic acceleration RMS. |
| Motion regime thresholds | $< 0.6 / 0.6\text{--}3.0 / > 3.0$ | Low / moderate / high-motion regimes. |
| Low-motion respiratory band | 0.038–1.00 Hz | Band-pass for low-motion recordings. |
| Moderate-motion respiratory band | 0.106–0.94 Hz | Band-pass for moderate-motion recordings. |
| High-motion respiratory band | 0.164–0.58 Hz | Band-pass for high-motion recordings. |
| Respiratory filter order | 8 (Butterworth) | Offline zero-phase band-pass filter. |
| RR window length (Rest/Exercise) | 39.34/37.85 s | Sliding window for RR estimation. |
| RR step size (Rest/Exercise) | 7.48/13.95 s | Step between consecutive windows. |
| Physiological RR range | 3–50 breaths/min | Validity range for retained windows. |

### S1.4 Respiratory Signal-Quality Score

The RR–SQS was computed after respiratory-rate extraction for every retained window. It combined three waveform descriptors: zero-crossing inflation, autocorrelation at the estimated respiratory lag, and interval regularity. Zero-crossing inflation penalised waveforms with more derivative zero crossings than expected from the estimated breathing rate:

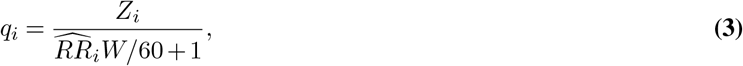

where *Z*_*i*_ is the number of derivative zero crossings and *W* is the window length. Interval regularity was

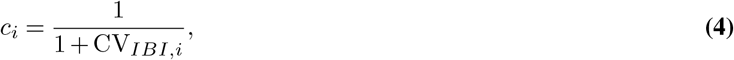

computed from peak-to-peak intervals when at least three peaks were available. Each component was globally rank-normalised onto [0, 1]; zero-crossing inflation was inverted, whereas autocorrelation and regularity were non-inverted. The composite score was

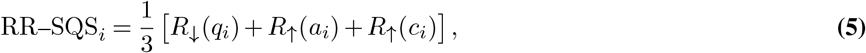

where *a*_*i*_ is the autocorrelation component, *R*_↓_ denotes inverted rank normalisation, and *R*_↑_ denotes non-inverted rank normalisation. Windows with RR–SQS below a set RR–SQS threshold or non-finite score were masked. Bounded interior gaps were linearly interpolated when at least two finite neighbours remained; leading and trailing low-quality windows stayed missing. Settings are summarised in Supplementary Table 3.

**Table 3.** Respiratory signal-quality-score settings.

| Parameter | Value | Description |
| --- | --- | --- |
| Scoring unit | RR analysis window | Same windows as RR estimation. |
| Components | Zero-crossing, autocorrelation, regularity | Waveform complexity, periodicity, interval regularity. |
| Normalisation | Global rank, $[0, 1]$ | Across all processed RR windows. |
| Composite | Mean of three components | Final RR–SQS. |
| Masking threshold | 0.2842 | Windows below threshold masked. |
| Interior-gap handling | Linear interpolation | Bounded gaps with at least two finite neighbours. |
| Edge-gap handling | Missing | Low-quality leading/trailing windows not interpolated. |

### S1.5 Form-Factor and Operating-Limit Analyses

The adhesive and magnet-through-textile recordings were processed independently with the same respiratory-rate pipeline. For each participant and task with paired valid data, magnet-minus-adhesive differences were computed for mean absolute error, bias, limits-of-agreement width, valid-window fraction, and percentage within 2 breaths/min. Negative differences on error metrics and positive differences on threshold-accuracy metrics indicate better magnet performance. Rest/controlled and exercise tasks were summarised separately.

Operating-limit analyses related respiratory-rate error to RR–SQS, motion intensity, reference respiratory rate, ventilation range, posture, form factor, and exercise stage. The primary visualisation plotted signed error against RR–SQS and marked the locked masking threshold. Recording-level summaries related median absolute error to within-recording accelerometer range and reference-rate range. Representative low-quality windows were inspected to identify dominant failure modes. These analyses were interpreted descriptively as operating-envelope summaries rather than as confirmatory hypothesis tests.

### S1.6 Oral/Nasal Classification

Controlled oral-only and nasal-only Phase A recordings were analysed as a secondary classification task. After task-onset trimming, the magnet-form-factor acoustic signal was segmented into 10 s windows with a 1 s step. The signal was median-centred, robustly clipped, notch filtered for power-line contamination, and transformed into log Hilbert envelopes in four acoustic bands spanning 2–120 Hz. Segment-level features were selected with an mRMR feature selector and classified with a robust-scaled shrinkage linear discriminant model. The primary endpoint used target-only participant-specific calibration with 20 s/class of oral and nasal data; calibration and held-out test segments were separated in time to limit dependence from overlapping windows. Balanced accuracy was primary. Sensitivity analyses tested zero-shot transfer, source-plus-target training, alternative calibration durations, and feature-family ablations. Design-relevant settings are listed in Supplementary Table 4.

**Table 4.**
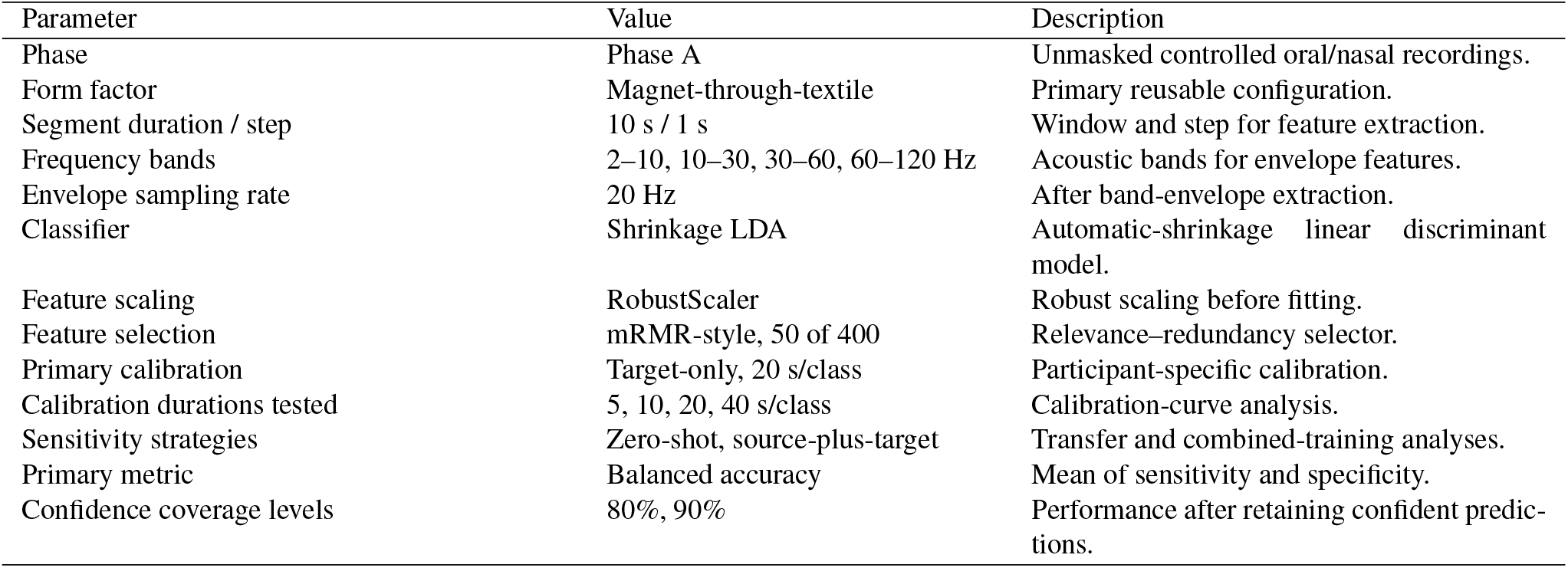
Design-relevant oral/nasal classification parameters. Full feature and selector settings are specified in the released configuration.

| Parameter | Value | Description |
| --- | --- | --- |
| Phase | Phase A | Unmasked controlled oral/nasal recordings. |
| Form factor | Magnet-through-textile | Primary reusable configuration. |
| Segment duration / step | 10 s / 1 s | Window and step for feature extraction. |
| Frequency bands | 2–10, 10–30, 30–60, 60–120 Hz | Acoustic bands for envelope features. |
| Envelope sampling rate | 20 Hz | After band-envelope extraction. |
| Classifier | Shrinkage LDA | Automatic-shrinkage linear discriminant model. |
| Feature scaling | RobustScaler | Robust scaling before fitting. |
| Feature selection | mRMR-style, 50 of 400 | Relevance–redundancy selector. |
| Primary calibration | Target-only, 20 s/class | Participant-specific calibration. |
| Calibration durations tested | 5, 10, 20, 40 s/class | Calibration-curve analysis. |
| Sensitivity strategies | Zero-shot, source-plus-target | Transfer and combined-training analyses. |
| Primary metric | Balanced accuracy | Mean of sensitivity and specificity. |
| Confidence coverage levels | 80%, 90% | Performance after retaining confident predictions. |

### S1.7 Sensitivity Analyses

Robustness checks were grouped around the predefined endpoints: reference-specific respiratory-rate agreement for spirometry, SLP, and paced targets; task-specific stratification by baseline, paced rate, posture, airway task, exercise stage, cool-down, and recovery; participant-level summaries to avoid treating overlapping windows as independent; respiratory-rate error stratified by RR–SQS, motion, and reference-rate range; paired-only form-factor comparison restricted to strata with valid data from both devices; oral/nasal performance across calibration durations; oral/nasal feature-family ablations; zero-shot oral/nasal transfer; and source-plus-target oral/nasal training. The primary interpretation rests on magnet-form-factor respiratory-rate agreement, paired form-factor comparison, and the locked calibrated oral/nasal endpoint; the remaining analyses define operating limits and exploratory signal content.

## S2 Supplementary Results

### S2.1 Demographic Associations

We explored whether participant-level signal quality or respiratory-rate error varied with BMI, age, body mass, height, or sex. Height was measured using a portable stadiometer (Seca 217; Seca GmbH & Co. KG, Hamburg, Germany), and body mass was measured using a mechanical flat scale (Seca 760; Seca GmbH & Co. KG, Hamburg, Germany). Continuous associations used Spearman’s rank correlation (*ρ*), with Pearson’s *r* reported as a sensitivity summary; sex differences used a Mann–Whitney *U* test. These analyses were exploratory. Ten uncorrected tests were performed, and individual *p*-values should therefore be interpreted descriptively (Supplementary Figure 5).

Median RR–SQS was lower in participants with higher BMI (*ρ* = −0.63, *p* = 0.003) and higher body mass (*ρ* = −0.51, *p* = 0.021). These associations are consistent with participant-level differences in sensor coupling, but body-composition data were not collected, so the mechanism cannot be determined from the present study. The reduction in signal quality did not translate into a comparable increase in respiratory-rate error: BMI was only weakly associated with MAE (*ρ* = 0.30, *p* = 0.19). This dissociation is consistent with the RR–SQS masking stage, which removes low-quality windows before endpoint agreement is estimated.

Age showed no robust monotonic association with signal quality or error. The Pearson correlation between age and MAE was significant (*r* = 0.52, *p* = 0.018), whereas the Spearman correlation was not (*ρ* = 0.31, *p* = 0.18), suggesting sensitivity to one or more high-leverage older participants. Height was not associated with either outcome (|*ρ*| ≤0.25, all *p* > 0.28). Signal quality was nominally lower and MAE nominally higher in female participants (median RR–SQS 0.439 versus 0.490; MAE 0.783 versus 0.685 breaths/min), but neither difference was significant (*p* = 0.74 and *p* = 0.20), and the small female subgroup limits interpretation. Overall, BMI was the only demographic variable with a clear association with signal quality, whereas respiratory-rate error remained weakly associated with the measured demographic variables.

### S2.2 Signal Quality Analysis

Median RR–SQS varied systematically across tasks and participants (Supplementary Figure 6). Speech and scripted-motion tasks showed consistently low signal quality, whereas paced-breathing tasks in Phases A and B showed high signal quality, consistent with their periodic, low-motion waveforms. Quiet breathing, posture tasks, and airway-mode tasks showed intermediate signal quality and stronger participant-to-participant variability.

RR–SQS also varied across participants independently of task. Several participants, including for example participants 8, 13, and 22, showed lower signal quality across multiple task groups. These participant-level differences are consistent with the demographic associations reported above, but may also reflect differences in sensor placement, garment fit, or mechanical coupling that the present design cannot disentangle.

### S2.3 Exploratory Waveform Descriptors

Because the processing pipeline segments individual breaths, the same respiratory waveform used for rate estimation also yields candidate breath-resolved descriptors: inhalation and exhalation duration, total breath duration and their ratios, breath-to-breath interval variability, and a peak-to-trough tidal-amplitude proxy (Supplementary Figure 7; representative recording from Stage 1 of the Bruce protocol). Panel (a) shows the breath-to-breath interval time series varying around its mean, and panel (b) shows the per-breath inhalation/exhalation segmentation and amplitude measurement on the processed waveform.

These descriptors are reported as exploratory capability only. The exercise and masked references used here (breath-by-breath spirometry) provide respiratory rate and ventilation but not the breath-resolved inspiratory and expiratory timing or a calibrated tidal-volume reference against which these quantities could be validated. Accuracy assessment of waveform timing and amplitude is therefore out of scope for the present study and is deferred to future work with phase- and volume-resolved references, as noted in the Discussion.

### S2.4 Usability Analysis

All 24 participants completed a post-session acceptability questionnaire covering device comfort, movement restriction, form-factor preference, and an adapted 10-item System Usability Scale (Supplementary Figure 8). Both Alveos One form factors were rated favourably for comfort, with most participants selecting one of the top two categories for the magnet configuration (20/24) and adhesive patch (18/24). Participants reported little movement restriction during exercise for either form factor, with 20/24 selecting the lowest category for both configurations.

When asked which form factor they would prefer for extended daily or overnight use, most participants preferred the magnet configuration (13/24, 54%) over the adhesive patch (4/24, 17%); seven participants expressed no preference (7/24, 29%). The adapted System Usability Scale yielded a median score of approximately 83/100. These questionnaire results support the practical acceptability of the reusable magnet configuration in a supervised laboratory session, while longer free-living studies will be needed to assess comfort, adherence, and skin tolerance during repeated or prolonged wear.

## Notes

https://github.com/alveoslabs/alveos-respiratory-validation

